# Early Loss of Deep Restorative Sleep and Auditory Stimulus Evoked 40-Hz activity of Hippocampal Parvalbumin Neurons in the APP/PS1 Mouse Model of Alzheimer’s Disease

**DOI:** 10.64898/2026.05.26.725476

**Authors:** Fumi Katsuki, James M. McNally, Dmitry Gerashchenko, David S. Uygun, Alyssa Tyler, John G. McCoy, James T. McKenna, Ritchie E. Brown

## Abstract

Sleep abnormalities and dysfunction of gamma band (30–80 Hz) activity generated by parvalbumin (PV) interneurons are early characteristics of Alzheimer’s disease (AD) which correlate with the severity of amyloid-β deposition (Aβ) and cognitive impairment. However, the timing of these alterations *in vivo* with respect to disease progression is unclear. Here, in longitudinal recordings from APP/PS1/PV-cre (AD mice) from 3–6 months, we found reduced sleep slow-wave power (0.5–4 Hz) in hippocampus and medial prefrontal cortex in AD mice as young as 3 months old, compared to non-AD (PV-cre) mice, well before overt pathology. This finding was primarily due to reductions in the NREM delta range (1.5–4 Hz), a hallmark of restorative functions of sleep. In contrast, beta (15–30 Hz) power linked to insomnia was significantly higher across all sleep-wake states. Loss of deep NREM sleep was not compensated by an increase in NREM sleep time, instead NREM sleep during the dark (active) phase was slightly but significantly lower in AD mice. 40-Hz auditory steady-state responses and associated evoked calcium responses of hippocampal PV neurons recorded using fiber photometry were also impaired by 3 months old. However, Y-maze performance in 3- and 6-month-old AD mice was not significantly different from non-AD mice. These results reveal reduced deep sleep and PV-associated 40-Hz activity as very early changes amenable to early intervention occurring prior to cognitive deficits. Furthermore, they establish APP/PS1 mice as a good model to causally test the relationship between sleep, PV neuronal activity and amyloid-mediated pathology.

## INTRODUCTION

Sleep abnormalities are one of the commonly reported phenotypes of Alzheimer’s disease (AD) linked to pathology and cognitive impairments (Guarnieri et al., 2012; Mander et al., 2015; Wang & Holtzman, 2020; Wennberg et al., 2017). The sleep abnormalities in AD include changes in overall sleep pattern such as decreased total sleep time and fragmentation of non-rapid eye movement (NREM) sleep and REM sleep (Bliwise et al., 1989; Bonanni et al., 2005; Duncan et al., 2022; Liguori et al., 2014; Lim et al., 2013; Mander et al., 2016; Vitiello et al., 1990). Notably, the sleep changes become apparent at the preclinical stage of AD and get worse over the course of the disease (Mander et al., 2016).

Studies in AD patients and animal models also show prominent impairments in sleep related brain oscillations including changes in slow-wave activity (0.5–4 Hz) (Byron et al., 2021; Katsuki et al., 2022). NREM slow-wave activity refers to neural oscillations with high amplitude and low frequency ranging from 0.5 to 4 Hz, which can be further subdivided into cortical slow oscillations (<1 Hz) and thalamocortical delta oscillations ranging between 1.5 and 4 Hz (Kim et al., 2019; Steriade, 2006; Steriade, Amzica, et al., 1993; Steriade, Contreras, et al., 1993; Steriade, Nunez, et al., 1993b, 1993a; Steriade & Timofeev, 2003; Uygun & Basheer, 2022).

Slow-wave activity defines the deeper stages of NREM sleep (Steriade, Amzica, et al., 1993; Steriade, Nunez, et al., 1993a, 1993b) and is linked to many of the restorative effects of sleep (Huber et al., 2000). Slow-wave activity, especially in the delta range, is considered a hallmark of sleep intensity since it is elevated during the recovery sleep following an acute sleep deprivation (Achermann et al., 2001).

Reduced NREM slow-wave activity has been observed in EEG recordings from midline frontal, central, and parietal regions in cognitively normal older adults with amyloid-β (Aβ) or tau pathology (Lucey et al., 2019a; Mander et al., 2013, 2015; Varga et al., 2016) and in patients with mild cognitive impairment (MCI) (Westerberg et al., 2012), suggesting that abnormal NREM slow-wave activity could emerge at preclinical or the early, asymptomatic stage of AD. These abnormalities and the severity of deposition of Aβ and cognitive impairment during later stages of the disease are correlated (Y. E. Ju et al., 2014; Mander et al., 2016). Furthermore, sleep abnormalities in AD are linked to impaired glymphatic waste clearance of Aβ and tau, which is normally facilitated during NREM sleep in both humans and rodents (Holth et al., 2019; Jessen et al., 2015; Y. E. S. Ju et al., 2017; Kang et al., 2009a; Mander et al., 2015; Morawska et al., 2016). Although these findings suggest potential links, the exact temporal and causal relationships between sleep abnormalities and increased Aβ and Tau burden in AD and their time course are still active and unresolved areas of investigation important for the development of early therapeutic interventions.

Abnormal fast cortical activity in the beta and gamma range generated by fast-spiking cortical interneurons which contain the calcium binding protein PV, is another early-stage characteristic of AD (Güntekin et al., 2013; Katsuki et al., 2022; Palop et al., 2007; Verret et al., 2012), contributing to AD-related cortical network dysfunction and progression of AD pathology (Palop & Mucke, 2010; Styr & Slutsky, 2018). Conversely, enhancement of 40-Hz activity has been proposed as a therapeutic strategy to delay progression (Adaikkan et al., 2017; Chan et al., 2025; Cimenser et al., 2021; Iaccarino et al., 2016; Martorell et al., 2019). However, how the ability to generate resonant gamma oscillations changes over time in AD models *in vivo* is unclear. A previous *in vitro* study in a rodent Aβ AD model showed that hippocampal (HPC) PV neurons in APP/PS1 mice are hyperexcitable at an early stage (3-4 months old) in disease progression, prior to changes in pyramidal neuronal excitability, and this was linked to abnormal neuronal network activity and memory impairment (Hijazi et al., 2019). Conversely, another *in vitro* study in 5XFAD mice reported reduced activity of HPC PV basket cells in early amyloid pathology (Caccavano et al., 2020). However, the direction and timing of changes in gamma band activity *in vivo* is unclear. Moreover sleep-wake states are not robustly conserved *in vitro*.

Sleep disruption and abnormal activity of PV neurons both occur early in AD and are suggested to be potential factors for accelerating AD pathogenesis. However, the timing and longitudinal progression of these two changes in AD is not fully characterized since most prior animal studies have used a cross-sectional design and focused on later stages of progression (Katsuki et al., 2022). In contrast, here we used longitudinal recordings beginning at the young adult stage to investigate whether changes in sleep oscillations and hippocampal PV activity occur prior to cognitive deficits and overt pathology in APP/PS1 mice, one of the best characterized Aβ AD mouse models with mutations in two human AD related genes, amyloid precursor protein (hAPP) and presenilin 1 (hPSEN1). We find that indeed, changes in sleep oscillations and hippocampal PV activity occur well before pathological changes and cognitive deficits are apparent.

## METHODS

### Animals

To target the hippocampal interneurons which express PV in AD mice, we crossed PV-Cre mice (B6.129P2-Pvalbtm1(cre)Arbr/J; Stock #: 017320; Jackson Laboratory, Bar Harbor, ME, USA) with APP/PS1 mice. In a subset of experiments PV-cre mice were crossed with another amyloid-over expressing strain, 5XFAD mice. The APP/PS1 mice used for this project, B6J;C3H-Tg (APPswe,PSEN1dE9) 85Dbo/Mmjax, RRID:MMRRC_034829-JAX, were obtained from the Mutant Mouse Resource and Research Center (MMRRC) at The Jackson Laboratory (Bar Harbor, ME, USA), an NIH-funded strain repository, and was donated to the MMRRC by David Borchelt, Ph.D., McKnight Brain Institute, University of Florida (Jankowsky et al., 2001).

APP/PS1 mice express a chimeric mouse/human amyloid precursor protein (Mo/HuAPP695swe) and a mutant human presenilin 1 (PS1-dE9). Forty percent of mice used in the current project were born from homozygous PV-Cre females crossed with hemizygous APP/PS1 males, and the rest were born from hemizygous APP/PS1 females crossed with homozygous PV-Cre males. The 5XFAD mice used for this research project, B6SJL-Tg (APPSwFlLon,PSEN1*M146L *L286V)6799Vas/Mmjax, RRID:MMRRC 034840-JAX, were obtained MMRRC at The Jackson Laboratory, an NIH-funded strain repository, and was donated to the MMRRC by Robert Vassar, Ph.D., Northwestern University. 5XFAD transgenic mice overexpress mutant human amyloid beta (A4) precursor protein 695 (APP) with the Swedish (K670N, M671L), Florida (I716V), and London (V717I) Familial Alzheimer’s Disease (FAD) mutations along with human presenilin 1 (PS1) harboring two FAD mutations, M146L and L286V. Sixty percent of mice were born from homozygous PV-Cre females crossed with hemizygous 5XFAD males, and the rest were born from hemizygous 5XFAD females crossed with homozygous PV-Cre males. Genotyping of crossed animals (APPPS1/PVCre or 5XFAD/PVCre) were performed by TransnetYX (Cordova, TN, USA) to identify if mice were APPsw/PSEN1 positive or negative. Littermate mice identified as APPsw/PSEN1 negative were used as non-AD control mice.

Additional non-littermate PV-Cre mice were also used as non-AD mice. The female-to-male ratios used in the experiments were 5:6 for APP/PS1 AD positive mice and 4:9 for non-AD mice, 1:1 for 5XFAD AD positive mice and 4:3 for non-AD mice. There were no obvious sex differences, thus data from both sexes were combined. Mice were housed at room temperature with 12:12 hour light/dark cycle, and food and water available *ad libitum*. All procedures were performed in accordance with National Institutes of Health guidelines and in compliance with the animal protocol approved by the Institutional Animal Care and Use Committee (IACUC) of the Veterans Administration Boston Healthcare System.

### Stereotaxic surgeries

All surgeries were performed at 2 months old under aseptic conditions in anesthetized mice using isoflurane (induction, 5%; maintenance, 1–2%). After surgery, analgesic (Meloxicam, 5mg/kg) was given intramuscularly or subcutaneously, again 22-24 hours later, and each mouse was housed in individual cages and allowed to recover from surgery for at least 2 weeks. LFP electrodes were implanted in mPFC (AP 1.9 mm, ML −0.45 mm, DV −1.7 mm) and HPC CA1 (AP −2.0 mm, ML −1.8 mm, DV −1.25 mm) (**Fig. 1A**). A reference screw and a ground electrode were placed above the cerebellum. Electromyography (EMG) electrodes were inserted in the nuchal muscle. Electrodes were then soldered to a headmount (Cat#: 8201, Pinnacle Technology Inc., Lawrence, KS, USA). The fiber-optic cannula for the fiber photometry recording (Mono fiber-optic cannula: MFC_200/250-0.66_8.0mm_MF1.25_FLT, Doric Lenses, Québec, QC, Canada) was implanted targeting HPC on the contralateral side to the LFP electrode (AP −2.0 mm, ML 1.8 mm, DV −1.25 mm) (**Fig. 1A**). The electrode, the headmount and the fiber-optic cannula were secured to the skull using dental cement (Fastray, Keystone Industries, Gibbstown, NJ, USA, or Ortho-Jet, Lang Dental Manufacturing Company, Wheeling, IL, USA).

**Fig 1.**
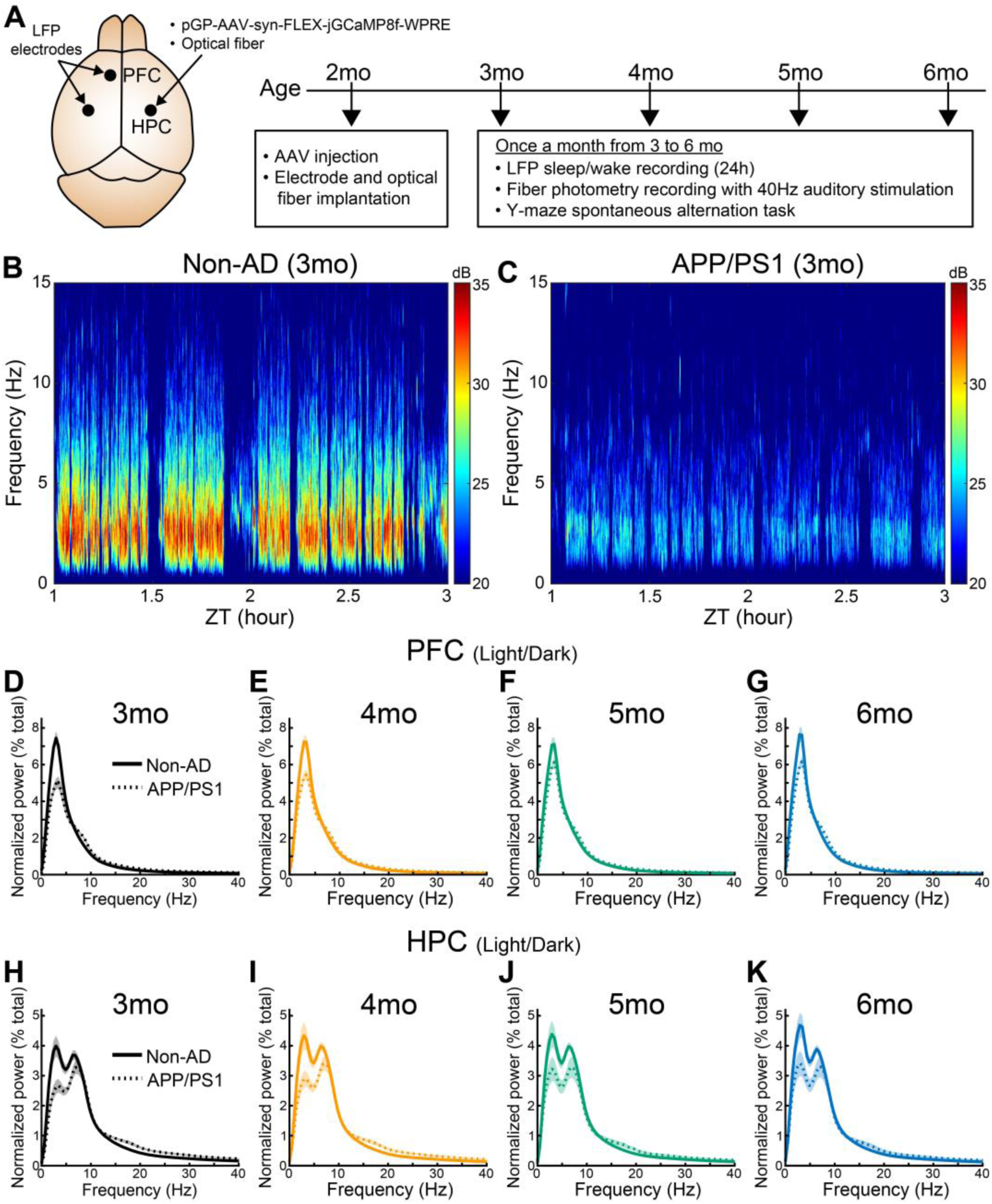
Longitudinal recordings revealed that spectral powers of brain oscillations were abnormal in APP/PS1 mice as early as 3 months old and remained abnormal until 6 months. (A) Local field potential (LFP) electrodes were implanted in frontal cortex (PFC) and hippocampus (HPC). AAV5-GCaMP8f injection was performed in HPC, along with an optical fiber implantation for fiber photometry recordings of calcium activity from PV neurons. The diagram shows the experimental timeline. (B-C) Representative spectrograms of a non-AD mouse (left) and an APP/PS1 mouse (right) at 3 months old. There was a striking reduction in power at the lower frequency range (<4 Hz) in the APP/PS1 mouse compared to the non-AD mouse. Data are plotted for zeitgeber time (ZT) 1–3 hour (light phase). (D-K) Averaged power spectral density (PSD) plots from PFC (D-G) and HPC (H-K) LFP recordings showed lower power in the slow-wave activity range (0.5–4Hz) in APP/PS1 mice (N=11) compared to non-AD mice (N=13) as early as 3 months of age which were largely maintained in the older animals. On the other hand, there were increases in high theta (7–9Hz) power in PFC and beta power in both areas in APP/PS1 mice. Figures were plotted with data including all sleep/wake states recorded from 24 h (both light and dark phase). Shaded areas represent SEM.

### *In vivo* 24h electrophysiological recordings and data analysis

#### LFP recordings

LFP recordings from mPFC and CA1 for a continuous 24-h were performed monthly at 3, 4, 5, and 6 months of age. Before beginning each experiment, animals were acclimatized to the recording chamber for two days. LFP/EMG signals (LFP: 2 KHz sampling, 0.5–650 Hz band pass filtered; EMG: 100Hz sampling, 1–100Hz band pass filtered) were recorded using Sirenia Acquisition (Pinnacle Technologies, Inc.). Data were then sleep scored and analyzed offline using MATLAB as described in the following section (R2019b and R2023b, MathWorks, Natick, MA, USA).

#### Sleep-wake Scoring

Sleep-wake scoring was performed using Sleep-Deep-Learner (Katsuki et al., 2024), our validated transfer learning-based automated sleep-wake scoring tool. Sleep-Deep-Learner first learns to score from the sample manual scores provided by the investigator for each file and scores the rest of the data on the file. Manual sleep-wake scoring was performed using Sirenia (Pinnacle Technologies, Inc., Lawrence, KS, USA) as we have previously described (Thankachan et al., 2019). Briefly, state was determined in 4s epochs using a combination of LFP/EMG measures. Wake was defined as epochs where LFP had desynchronized, low amplitude signals with muscle tone in the EMG signals (both tonic and phasic). NREM was defined as epochs where LFP had slow, synchronized, high-amplitude signals, paired with low amplitude EMG signals. REM sleep epochs were characterized by LFP signals with prominent theta activity (5–9 Hz) with essentially flat EMG signals occasionally contaminated by a pulse not indicative of skeletal muscle tone. The epochs with artifacts (e.g. due to crosstalk between LFP and EMG signals or brief water bottle contacts on the implant) were labelled simply as artifacts or artifacts with a specific state where the state was still identifiable. For sleep architecture analysis, such as the proportions of states or bouts, epochs including artifacts with identified state were used. The epochs with artifacts were excluded for analysis that used actual LFP signals including spectral power analysis. For data files (8 out of 96 files) which were flagged for low performance by Sleep-Deep-Learner (Katsuki et al., 2024), entire sleep-wake scoring was performed manually. Sleep-Deep-Learner and subsequent analyses of data after sleep-wake scoring were performed in MATLAB (Mathworks, R2019b, R2023b, Natick, MA, USA).

#### Spectral analysis

Spectral power analysis was performed using a MATLAB function “pwelch” with 2-second Hanning window, overlap 50%. Normalized power was determined by the percentage of each frequency bin per state of the total summed power of the whole 24h signal, as in prior work (Uygun et al., 2022). Power within defined frequency bands were computed for slow-wave (0.5–4 Hz), slow-oscillation (0.5–1 Hz), delta (1.5–4 Hz), low theta (4–7 Hz), high theta (7–9 Hz), sigma (10–15 Hz), beta (15–30 Hz), and gamma (30–80 Hz) frequency ranges.

#### Spindle detection procedure

Sleep spindles were detected post hoc using an automated algorithm developed and validated in house previously (Uygun et al., 2019), which utilized custom MATLAB scripts, and involved the following steps. First, raw EEG data was band-pass filtered across the spindle frequency range (10-15 Hz, Butterworth Filter). The root mean squared (RMS) power was calculated and then smoothed, providing an upper envelope of the data. The RMS data was then exponentially transformed, to accentuate spindle-generated signals over baseline. Putative spindle peaks were then identified in transformed data via crossing of an upper-threshold value, based on the background spindle frequency activity for each mouse (3.5X mean). Spindle duration was then defined by summing the time before and after the detected peak till the data drops below a lower-threshold value (1.2X mean). Events that were less than 0.5s were excluded from detection. Additionally, to account for trains of spindles that could not reliably be resolved, a minimum inter-event interval criteria was set at 0.1s. If juxtaposing spindles failed to meet this criterion, they were counted as a single event.

### Fiber photometry recordings and LFP recordings with 40-Hz auditory stimulation

#### Viral vector

Fiber photometry recordings were performed to capture the intracellular calcium population activity of the PV interneuron population in HPC CA1. We first injected each animal with 500 nL of adeno-associated virus (AAV) of pGP-AAV-syn-FLEX-jGCaMP8f-WPRE (Catalog #: 162379-AAV5, Addgene, Watertown, MA, USA) in HPC CA1 (AP −2.0 mm, ML 1.8 mm, DV −1.25 mm) (**Fig. 1A**) at 0.01 µl/minute using a 5 or 10 μl Hamilton Syringe and Legato 130 syringe pump (KD scientific, Holliston, MA, USA). The fiber-optic cannula was implanted at the injection site. There were 3–4 weeks after AAV injection before the beginning of experiments to allow sufficient transduction.

#### Fiber photometry recordings and analysis with 40-Hz auditory stimulation

Fiber photometry recording was performed as follows using a system from Doric Lenses (FPS_2S_GFP/ RFP, Doric Lenses Inc., Québec, Canada). Calcium-dependent fluorescence was excited by the 470 nm LED. The LED was coupled to a fiber optic mini-cube containing appropriate GFP excitation filters. Light was delivered to the animal via a fiber optic patch cable connected to an implanted fiber-optic cannula, in the targeting brain area. The emitted fluorescence was collected by the same fiber and directed back through a filter cube to an incorporated photodetector in the photometry recording device. The signals were then amplified via fluorescence detector amplifiers, demodulated via software-based lock-in approach.

Fiber photometry recordings and LFP recordings were performed simultaneously in the same mice monthly from 3 to 6 months of age while 40-Hz auditory stimulation was presented. The recordings were performed between ZT 6–9. This was done in addition to the sleep/wake recordings. The 40-Hz auditory stimulation (10 ms sound pulse duration) was on for 2 s followed by a 4 s of intertrial-interval and repeated for 150 trials (Spike2, Cambridge Electronic Design Limited, Milton, UK). The sound frequency tone for the auditory stimulation was at 10 kHz (Model 4052 Function Generator, B&K Precision, Yorba Linda, CA, USA), and the volume was set at 85 dB.

The demodulated fiber photometry signals were processed using MATLAB for further analysis. The signals were first detrended and adjusted with moving average baseline to obtain ΔF/F. The ΔF/F traces were normalized within each trial (z-score) aligned at the auditory stimulus onset. To assess PV neuronal responses to the 40-Hz auditory stimulation, we computed the areas under curve from 0.5 s to 2.5s of auditory stimulus onset where sustained PV activity was observed and compared them between APP/PS1 mice and non-AD mice for each month.

#### LFP recordings and analysis with 40-Hz auditory stimulation

Simultaneously with the fiber photometry recordings, we performed LFP recordings to track changes in brain oscillations during 40-Hz auditory stimulus presentation described above. LFP data were collected from mPFC and CA1. In order to assess the steady-state response during the auditory stimulation, first, the absolute power of background 40-Hz (35–45 Hz) activity was calculated from the pre-stimulation time (-2 to -0.2 s), then the absolute value of evoked 40 Hz power was calculated from the auditory stimulation time (0.2 to 4 s), excluding the period where large transient response was observed at around the onset of auditory stimulation (McNally et al., 2021). The steady-state response was computed as the percentage changes of 40-Hz power from pre-stimulation time to the stimulation time.

### Behavioral task

#### Y-maze spontaneous alternation task

Working-memory deficits are common in AD patients exhibiting abnormalities in brain regions including mPFC and HPC with high levels of Aβ deposition. Thus, spontaneous Y-maze alternation task which measures spatial working-memory and depends on the brain regions including mPFC and HPC (Lalonde, 2002) is suitable to assess cognitive aspects of AD progression. The task relies on mouse’s natural tendency to explore novel environment (Dudchenko, 2004) and does not require food/water deprivation or aversive reinforcement. Previous studies from other groups demonstrated impairments in this task at around 8 months old in APP/PS1 mice (Devi & Ohno, 2012; Shukla et al., 2013; Webster et al., 2014). Each mouse was placed in the center intersection of Y-maze and allowed to explore freely through the maze for 8 min. A EthoVisionXT14 video tracking system (Noldus Information Technology Inc., Leesburg, VA, USA) was used to track animals. The mice were tested with Y-maze task at 3, 4, 5, and 6 months of age within 1 week after each 24h LFP recording. In order to provide visual cues, different wallpaper inserts with black and white shapes/patterns were placed at10cm from the entrance of each arm of the Y-maze (Stoelting Co., Wood Dale, IL, USA). Each arm was equally illuminated at 30-35 lux. The combinations of shapes of wallpaper were changed each month. Percentage alternation of arm visits was computed as: number of triads containing entries into all three arms divided by the maximum possible alternations (the total number of arms entered minus 2) x 100 (Kimura et al., 2010; Oakley et al., 2006).

### Histology

After the experiments were completed, mice were perfused transcardially using phosphate buffered saline (PBS; 12–20 ml), then 4% paraformaldehyde (12 ml). Brains were post-fixed for 1–2 days in 4% paraformaldehyde and transferred to 30% sucrose in PBS. Coronal slices were cut (20–40 µm-thickness) with a microtome (Leica Biosystems, Illinois, United States) and stored in PBS at 4°C. Locations of optic fibers for fiber photometry in HPC and LFP electrodes in mPFC and HPC were confirmed using cresyl violet staining (**Supplemental Fig. 1**). Colocalization of PV neurons (anti-PV antibody, Cat# AF-5058, RnD Systems, 1:150) and GCaMP were validated with immunolabeling. The sections were mounted on the slides with coverslips secured by Vectashield Hard Set mounting medium (Part # H-1400, Vector Laboratories, Newark, CA, USA). Micrographs were acquired by Akoya Vectra Polaris Slide Scanner (Akoya Biosciences, Marlborough, MA, USA).

### Statistical analysis

All statistical analyses were performed in MATLAB. All data are presented as mean ± standard error. Paired comparisons of means were evaluated using Student’s t-test with significance set at p < 0.05. Multiple mean comparisons were performed using repeated measures ANOVA with Tukey-Kramer post-hoc analysis for pairwise comparisons.

## RESULTS

### Spectral power at low frequencies were lower in AD mice during NREM sleep in PFC and HPC at early ages

Given findings of reduced NREM slow wave activity in AD and MCI patients (Lucey et al., 2019b; Mander et al., 2015; Westerberg et al., 2012), we closely examined whether there are any changes in brain oscillations related to sleep/wake states in AD mice. Our LFP recordings were performed in both PFC and HPC monthly from 3 to 6 months old as shown in the experimental timeline (**Fig. 1A**). As representative plots of spectrograms show, APP/PS1 mice showed a dramatic reduction in spectral power in the low frequency range (< 4 Hz) compared to the non-AD mice (**Fig. 1B&C**). To quantitatively examine changes in the brain electrical oscillations, we first computed spectral power with data obtained during both light and dark phase (24 h recordings) regardless of sleep/wake states (**Fig. 1D-K**). Power at low frequencies (< 4 Hz) was lower in APP/PS1 mice in both PFC and HPC compared to non-AD mice. On the contrary, increased power in higher frequency ranges, including theta (7–9 Hz) and beta (15–30 Hz) were observed in APP/PS1 mice compared to non-AD mice (**Fig. 1D-K**). These differences were observed as early as 3 months old and were largely maintained until 6 months.

### Delta power was lower in APP/PS1 mice in PFC and HPC

We next looked at brain oscillatory changes during NREM sleep when slow-wave activity is most prominent (**Fig. 2A&E**). There was a large reduction in NREM slow-wave activity power (0.5–4Hz) in APP/PS1 mice as early as 3 months old compared to the non-AD mice both in PFC (**Fig. 2B**, % reduction at 3mo: 29.33%, 4mo: 25.40%, 5mo: 13.56%, 6mo: 20.90%) and HPC (**Fig. 2F**, % reduction at 3mo: 35.47%, 4mo: 34.53%, 5mo: 27.32%, 6mo: 25.44%). There was a statistically significant difference in genotypes on NREM slow-wave activity power in PFC data (F1,22=11.26, p=0.0029) and HPC data (F1,22=4.47, p=0.0461). The post-hoc pairwise comparisons indicated that slow-wave power was significantly lower in the AD group compared to non-AD group at 3, 4, and 6 months in PFC (3, 4, 5, 6 months, p=0.0032, p=0.00095, p=0.06, p=0.016, respectively) and at 3 and 4 months in HPC (3 months: p=0.018, 4 months: p=0.025). We further divided the slow-wave frequency into very slow side of frequency range (**Fig. 2C&G**, 0.5–1 Hz, i.e. the slow-oscillation) and the faster side of frequency range (**Fig. 2D&H**, 1.5–4 Hz, i.e. delta). The difference in slow oscillation power (0.5–1 Hz) between APP/PS1 mice and non-AD mice was not significant either in PFC or HPC (**Fig. 2C&G**). In contrast, delta power (1.5–4 Hz) was significantly lower in APP/PS1 mice compared to non-AD mice in both PFC (**Fig. 2D**, F1,22=12.19, p=0.0021, % reduction at 3mo: 30.87%, 4mo: 26.74%, 5mo: 14.79%, 6mo: 21.15%) and HPC (**Fig. 2H**, F1,22=4.79, p= 0.0395, % reduction at 3mo: 35.15%, 4mo: 34.94%, 5mo: 28.10%, 6mo: 26.37%). Lower delta power during NREM sleep in APP/PS1 was consistently found from 3 to 6 months old in the same mice, especially in PFC (**Fig. 2D**, 3, 4, 5, 6 months, p=0.0022, p=0.00061, p=0.048, p=0.015, respectively). Consistent with this APP/PS1 data, our smaller sample of data with 5XFAD mice, another commonly used Aβ mouse model, also showed lower slow-wave activity compared to non-AD mice starting around 3 months old and maintained at 6 months old (**Supplemental Fig. 2**).

**Fig 2.**
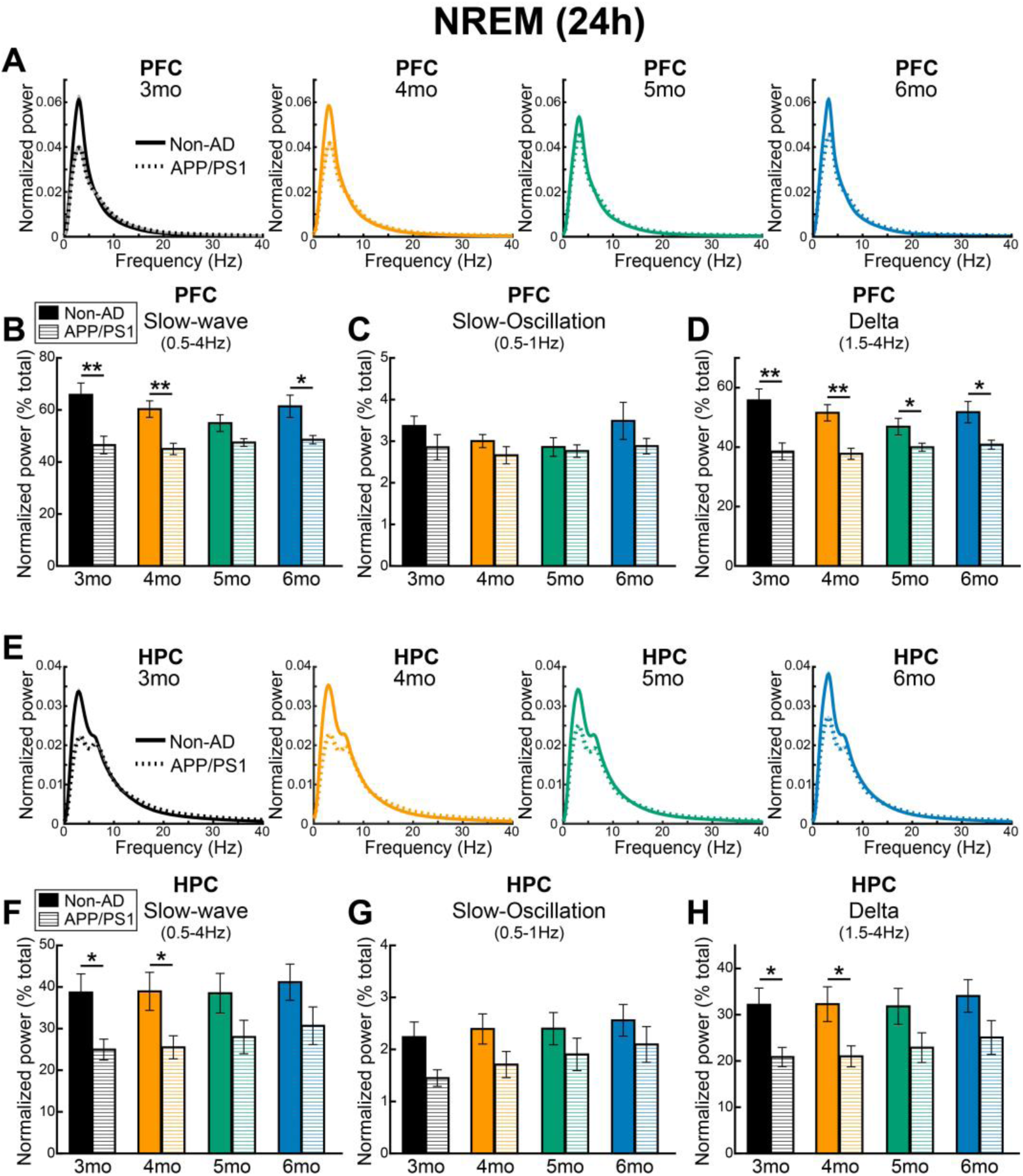
Slow-wave activity during NREM sleep was lower in APP/PS1 mice as early as 3 months old. **(A&E)** Averaged power spectral density (PSD) plots from PFC (**A**) and HPC (**E**). Analyses were focused on NREM sleep data from 24h recordings. **(B&F)** Large reduction in NREM slow-wave activity power (0.5–4Hz) in APP/PS1 mice (N=11) as early as 3 months old compared to non-AD mice (N=13) both in PFC and HPC. **(C-D&G-H)** When the slow-wave frequency was further categorized into very slow side of frequency range (**C&G**, 0.5–1Hz, slow-oscillation) and faster side of frequency range (**D&H**, 1.5–4Hz, delta), it became apparent that the reduction in slow-wave activity was mainly due to reductions in power in the delta range, while slow-oscillation changes were not significant in both PFC and HPC. Figures were plotted with data from NREM sleep state (both light and dark phase included). In older mice NREM delta activity in AD mice was also generally lower than in non-AD mice but did not reach significance in the hippocampus at 5 and 6 months old. Error bars represent SEM. Repeated measures ANOVAs with post-hoc Tukey-Kramer test was performed (*p<0.05, **p<0.01).

**Fig 3.**
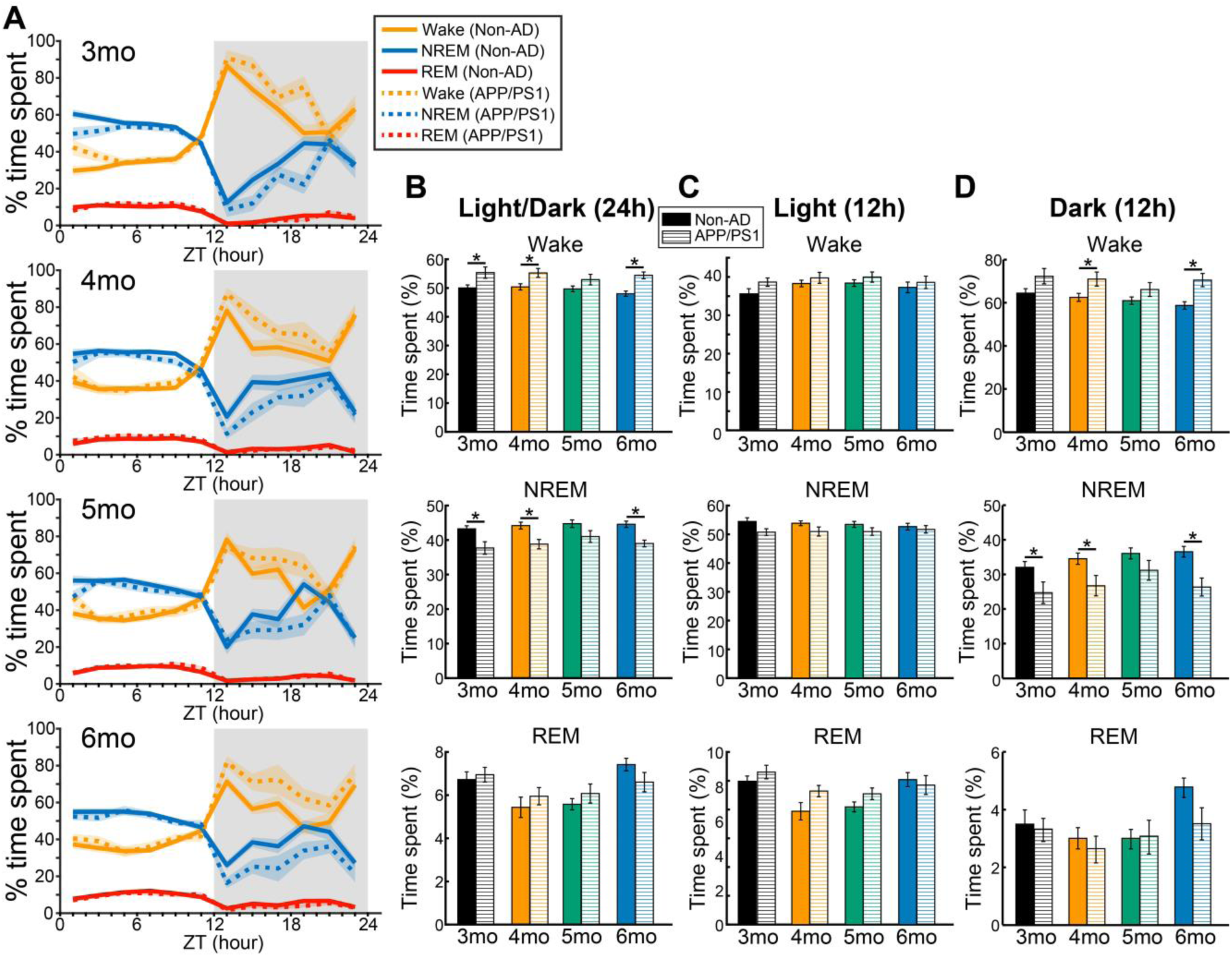
APP/PS1 mice tend to sleep less during the dark (active) phase. **(A)** Average percentage time spent in wake (orange), NREM sleep (blue) and REM sleep (red) are plotted over 24 h for non-AD (non-AD, N=13, solid lines) and APP/PS1 groups (N=11, dotted lines). **(B)** Over 24 hr, APP/PS1 mice spend less time in NREM sleep and more time in wakefulness compared to non-AD mice (significant at 3, 4 and 6 months). Time spent in REM sleep was not significantly changed. **(C)** In the light phase there were no significant changes in the percentage time spent in different behavioral states at any age (3-6 months). **(D)** In the dark phase wakefulness was generally higher in AD mice (significant at 4 and 6 months). Sleep scoring was performed based on the prefrontal LFP signal during the 24 h recordings. Repeated measures ANOVAs with post-hoc Tukey-Kramer test was performed (*p<0.05, **p<0.01). Shaded areas and error bars represent SEM.

**Fig 4.**
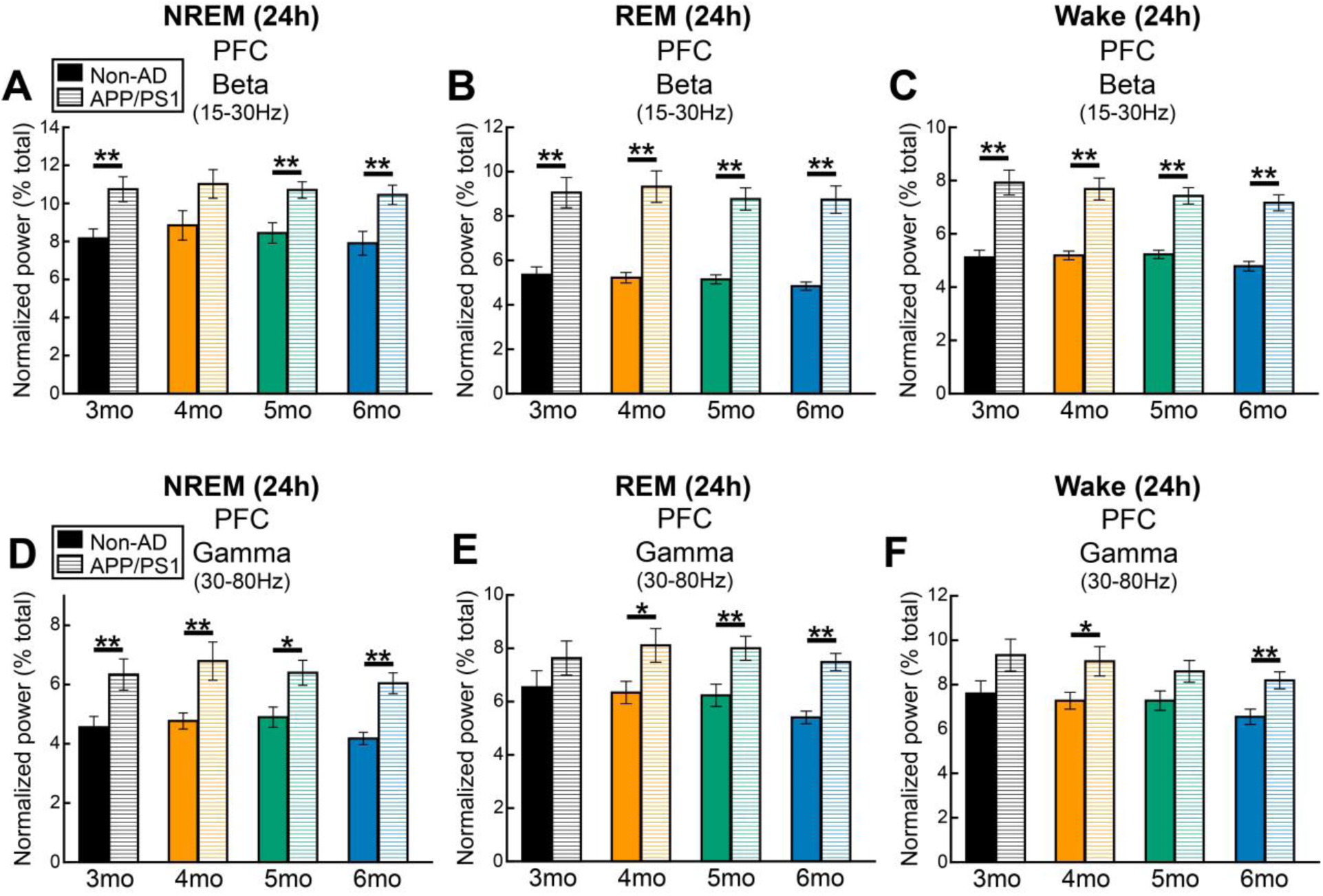
Beta and gamma power was higher in PFC of APP/PS1 mice across behavioral states as early as 3 months old. **(A-C)** Beta (15–30Hz) power was significantly higher in PFC of APP/PS1 mice (N=11) compared to non-AD mice (N=13). This increase was observed consistently at all ages during NREM sleep (**A**), REM sleep (**B**) and wakefulness (**C**). **(D-F)** Gamma power was significantly higher in APP/PS1 mice than the non-AD mice especially during NREM sleep (**D**) and REM sleep (**E**). The difference was less consistent during wakefulness (**F**). Both light and dark phases were included in the data. Error bars represent SEM. Repeated measures ANOVAs with post-hoc Tukey-Kramer test was performed (*p<0.05, **p<0.01).

**Fig 5.**
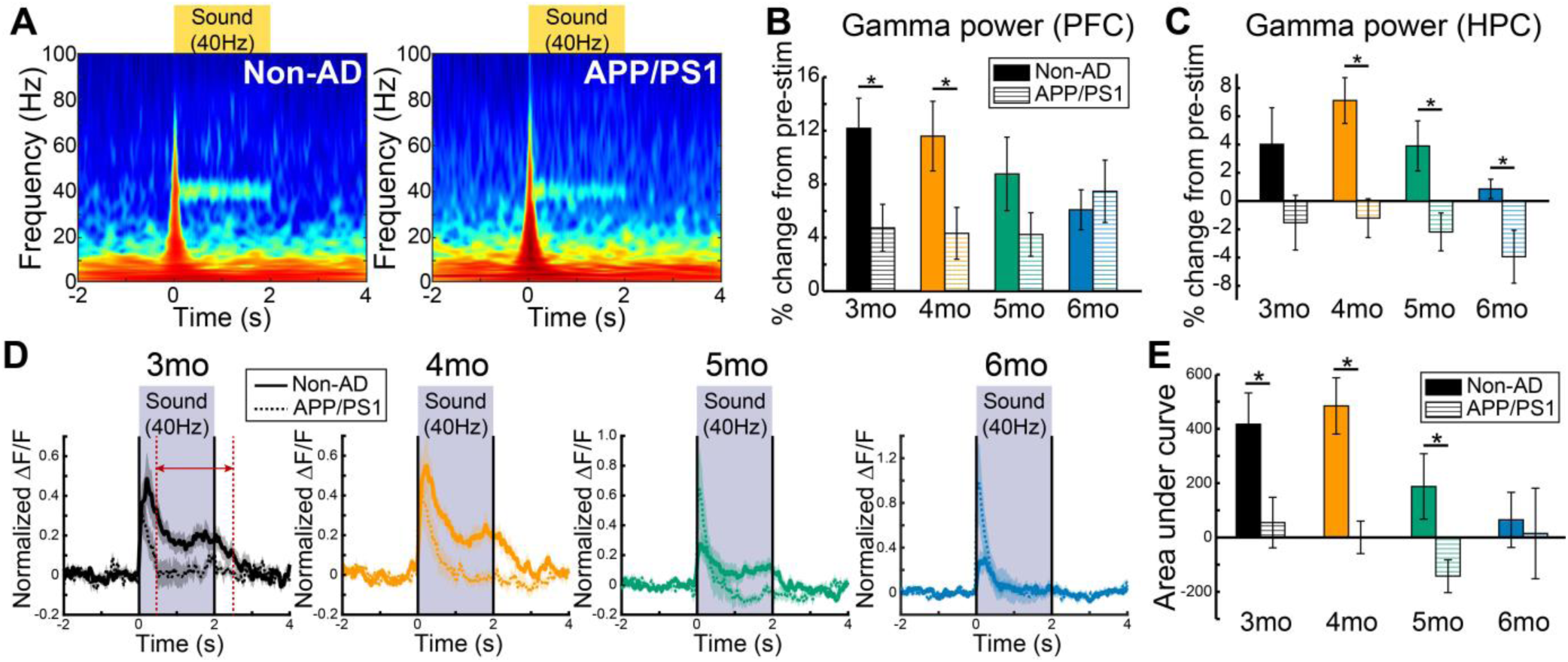
40-Hz electrical auditory steady-state response (ASSR) and hippocampal PV calcium response were lower in APP/PS1 mice. LFP recordings from mPFC and HPC and fiber photometry recordings of HPC PV neurons were performed simultaneously while 40-Hz auditory stimulation (2s-ON, 4s-OFF, 10 kHz at 85 dB, 150 trials) was presented to the mice. Recordings were repeated at 3, 4, 5, and 6 months old in the same mice. **(A)** Representative wavelet transformed LFP signals from 4 months old aligned at onset of 40-Hz auditory stimulation. A robust cortical evoked response at 40 Hz during the sound presentation can be observed in non-AD mouse (left), which was also seen in the APP/PS1 mouse but less robust (right). **(B&C)** Gamma power (35–45 Hz) during the time of 40-Hz auditory stimulation (0.2 to 4 s) was computed, then percentage changes from pre-stimulation time (-2 to -0.2 s) between non-AD (N=10) and APP/PS1 (N=9) groups were compared. Gamma power was significantly lower in APP/PS1 mice at 3 and 4 months old in PFC (**B**), and 4–6 months old in HPC (**C**) compared to non-AD mice. **(D)** Fiber photometry recordings of calcium activity from hippocampal PV neurons showed lower response in APP/PS1 mice as early as 3mo while 40-Hz auditory stimulation was presented. **(E)** Area under curves were computed using data from 0.5–2.5s of auditory stimulus onset (time between red dotted lines in Fig. D) to assess difference in PV neuronal response to the 40-Hz auditory stimulation. Significant differences were observed between non-AD and APP/PS1 groups at 3 and 4 months old. Shaded areas and error bars represent SEM. Statistical significance was tested with repeated measures ANOVAs with post-hoc Tukey-Kramer test (*p<0.05, **p<0.01).

### APP/PS1 mice sleep less during the active (dark) phase

Given their lower intensity of NREM sleep assessed by slow-wave power it might be expected that AD mice might need to sleep more to achieve the restorative effects of sleep. However, sleep scoring based on the prefrontal LFP signal during the 24 h recordings (**Fig. 3A**) revealed that APP/PS1 mice (N=11) spent significantly more time in wakefulness and at the expense of NREM sleep, compared to the healthy control (non-AD) mice (N=13) (**Fig. 3B**). A repeated-measures ANOVA was performed to compare the effect of genotypes on the percent time spent in each state. The percentage decrease in NREM sleep time in APP/PS1 mice compared to non-AD mice at each month were 12.86%, 12.18%, 8.34%, and 12.48% (**Fig. 3B middle**, 3, 4, 5, 6 months, respectively). In contrast, % increase in wakefulness in APP/PS1 mice compared to non-AD mice were, 10.65%, 9.64%, 6.52%, and 13.26% (**Fig. 3B top**, 3, 4, 5, 6 months, respectively). There was a statistically significant difference in genotypes on NREM sleep time (F1,22=12.94, p=0.0016). Tukey-Kramer post-hoc pairwise comparisons indicated that time spent in NREM sleep was significantly lower in APP/PS1 mice compared to non-AD group at 3, 4, and 6 months (3, 4, 5, 6 months, p= 0.0082, p= 0.0034, p= 0.07, p= 0.00043, respectively). On the other hand, time spent in wakefulness was significantly higher in APP/PS1 (F1,22=10.66, p=0.0035, post-hoc test: 3, 4, 5, 6 months, p= 0.020, p= 0.015, p= 0.12, p= 0.00027, respectively). Next, we looked at light phase (inactive phase for mice) and dark phase (active phase) separately. There was no significant difference between non-AD and APP/PS1 mice during the light phase in any state (**Fig. 3C**). During the dark (active) phase, % decrease in NREM sleep time in APP/PS1 mice were, 23.18%, 22.80%, 13.84%, and 28.07% (**Fig. 3D middle**, 3, 4, 5, 6 months, respectively) compared to non-AD mice. In contrast, % increase in wakefulness in APP/PS1 mice were 11.81%, 13.19%, 8.04%, and 19.60% (**Fig. 3D top,** 3, 4, 5, 6 months, respectively). There was a statistically significant difference in genotypes (F1,22=7.69, p= 0.011). The post-hoc pairwise comparisons indicated that time spent in NREM sleep was significantly lower in APP/PS1 mice compared to non-AD group at 3, 4, and 6 months (3, 4, 5, 6 months, p= 0.039, p= 0.023, p= 0.12, p= 0.0019, respectively). Conversely, time spent in wakefulness was significantly higher in APP/PS1 mice at 4 and 6 months (F1,22= 6.77, p= 0.016, post-hoc test: 3, 4, 5, 6 months, p= 0.063, p= 0.032, p= 0.17, p= 0.0025, respectively). There was no significant difference in REM sleep time (**Fig. 3B-D bottom**). The results showed that sleep patterns are already different in APP/PS1 mice as early as 3 months of age which is before brain-wide accumulation of Aβ (Garcia-Alloza et al., 2006; Jankowsky et al., 2004).

Furthermore, deficits in NREM slow-wave activity linked to restorative functions of NREM sleep are not compensated for by increases in NREM sleep time.

### Beta and gamma power was higher in APP/PS1 mice in PFC and HPC

Increased EEG beta activity across states and during sleep is associated with both primary insomnia (Spiegelhalder et al., 2012) and insomnia in Parkinson’s disease patients (Mizrahi-Kliger et al., 2020; Verma et al., 2024) as well as with diminished glymphatic function (Dagum et al., 2025; Hablitz et al., 2019). Thus, we next closely examined power in beta (15–30 Hz) frequency range and compared between APP/PS1 and non-AD mice during 24 h of recordings. Beta power was significantly higher in PFC of APP/PS1 mice compared to the non-AD mice when all sleep/wake states were included (Percentage increase at 3mo: 39.88%, 4mo: 36.21%, 5mo: 33.73%, 6mo: 39.07%, F1,22=24.74, p=0.000056). Post-hoc pairwise comparisons indicated that beta power was significantly and consistently higher in the AD group compared to non-AD group at all time points (3, 4, 5, 6 months, p=0.0022, p=0.00028, p=0.000061, p=0.00012, respectively). Higher beta power in the APP/PS1 group was also found in the HPC (Percentage increase at 3mo: 40.65%, 4mo: 42.24%, 5mo: 31.07%, 6mo: 29.42%, F1,2 =5.27, p=0.032), although pairwise significance was found only at 3 and 4 months (3 months: p=0.0040, 4 months: p=0.013). We further analyzed the data separately for each state (**Fig. 4A-C**). We found that beta power was significantly higher in APP/PS1 mice compared to non-AD mice, during wakefulness (**Fig. 4C**, PFC: Percentage increase 3mo: 54.99%, 4mo: 47.98%, 5mo: 42.07%, 6mo: 49.79%, F1,22=44.34, p=0.0000011, HPC: % increase 3mo: 56.65%, 4mo: 57.06%, 5mo: 46.41%, 6mo: 44.52%, F1,22=8.25, p=0.0089) and REM sleep (**Fig. 4B**, PFC: Percentage increase 3mo: 69.03%, 4mo: 78.74%, 5mo: 70.29%, 6mo: 80.79%, F1,22=43.96, p=0.0000011, HPC: % increase 3mo: 45.34%, 4mo: 43.01%, 5mo: 32.91%, 6mo: 37.98%, F1,22=3.97, p=0.059, figure for HPC not shown), more so than during NREM sleep. Beta power was also higher in APP/PS1 during NREM sleep, however, the magnitude of the increase was less than that of wakefulness and REM sleep (**Fig. 4A**, PFC: Percentage increase 3mo: 31.62%, 4mo: 24.66%, 5mo: 26.78%, 6mo: 32.15%, F1,22=9.15, p=0.0062, HPC: % increase 3mo: 24.43%, 4mo: 26.07%, 5mo: 17.19%, 6mo: 14.32%, F1,22=2.80, p=0.11). Higher beta activity was also observed in our PFC and HPC data obtained from the 5XFAD mice (**Supplemental Fig. 2**).

Dysfunction of gamma band (30–80 Hz) activity is another well-reported characteristic of AD (Goutagny et al., 2013; Iaccarino et al., 2016; Klein et al., 2016; Traikapi & Konstantinou, 2021). We found gamma power in PFC was significantly higher in APP/PS1 mice than non-AD mice especially during NREM sleep (**Fig. 4D**, Percentage increase 3mo: 39.04%, 4mo: 42.35%, 5mo: 30.61%, 6mo: 44.50%, F1,22= 11.55, p=0.0026, post-hoc test: 3, 4, 5, 6 months, p= 0.0094, p= 0.0059, p= 0.011, p= 0.00010, respectively) and REM sleep (**Fig. 4E**, % increase 3mo: 16.82%, 4mo: 28.08%, 5mo: 28.57%, 6mo: 38.70%, F1,22= 7.89, p= 0.01, post-hoc test: 3, 4, 5, 6 months, p= 0.23, p= 0.025, p= 0.0087, p= 0.000025, respectively). The difference was less consistent during wakefulness (**Fig. 4F**, Percentage increase 3mo: 22.76%, 4mo: 24.48%, 5mo: 17.99%, 6mo: 25.04%, F1,22= 6.09, p= 0.02, post-hoc test: 3, 4, 5, 6 months, p= 0.069, p= 0.025, p= 0.056, p= 0.004, respectively).

### Brain oscillation changes found in other frequency ranges in APP/PS1 mice in PFC

We also found brain oscillation changes in other frequency ranges including higher theta (7–9 Hz) and sigma (10–15 Hz). Powers in these frequency ranges were higher in APP/PS1 mice as early as 3 months old, however, differences were more state dependent than delta or beta frequencies (**Supplemental Fig. 3**). Spectral power around high theta range (7–9 Hz) was significantly higher in APP/PS1 mice compared to non-AD mice in PFC especially during REM sleep (**Supplemental Fig. 3C**, percentage increase 3mo: 57.95%, 4mo: 52.89%, 5mo: 43.44%, 6mo: 60.07%, F1,22= 17.42, p= 0.00039, post-hoc test: 3, 4, 5, 6 months, p= 0.00024, p= 0.0065, p= 0.0019, p= 0.000086, respectively) and wakefulness (**Supplemental Fig. 3E**, percentage increase 3mo: 36.73%, 4mo: 21.94%, 5mo: 16.67%, 6mo: 30.00%, F1,22= 15.09, p= 0.00080, post-hoc test: 3, 4, 5, 6 months, p= 0.000050, p= 0.021, p= 0.016, p= 0.00052, respectively) which was observed from 3 to 6 months old. Sigma power (10-15 Hz) was also higher in APP/PS1 mice compared to non-AD mice in PFC during REM sleep (**Supplemental Fig. 3D**, percentage increase 3mo: 65.13%, 4mo: 73.43%, 5mo: 52.26%, 6mo: 64.27%, F1,22= 27.34, p= 0.000030, post-hoc test: 3, 4, 5, 6 months, p= 0.00049, p= 0.00021, p= 0.00033, p= 0.000012, respectively) and wakefulness (**Supplemental Fig. 3F**, % increase 3mo: 39.77%, 4mo: 25.32%, 5mo: 18.86%, 6mo: 26.74%, F1,22= 26.69, p= 0.000035, post-hoc test: 3, 4, 5, 6 months, p= 0.000048, p= 0.00074, p= 0.00062, p= 0.00031, respectively), but not during NREM sleep (**Supplemental Fig. 3B**, % increase 3mo: 18.07%, 4mo: 7.92%, 5mo: 11.24%, 6mo: 14.37%). There was no statistically significant difference in NREM sleep spindle density (spindles/NREM min) between non-AD mice and APP/PS1 mice (non-AD vs. APP/PS1, 3mo: 5.66 ± 0.14 vs. 5.32 ± 0.11, 4mo: 5.52 ± 0.13 vs. 5.46 ± 0.19, 5mo: 5.54 ± 0.14 vs. 5.28 ± 0.12 6mo: 5.50 ± 0.12 vs. 5.34 ± 0.14 spindles/NREM min).

### Gamma power was lower during 40-Hz auditory stimulation in AD mice even at 3months old

40-Hz auditory and/or visual stimulation has been suggested as a potential AD therapy (Cimenser et al., 2021; Iaccarino et al., 2016). However, increases in broad-band gamma-band cortical activity, as we observed here, can cause impairments in the ability of cortical circuits to generate resonant gamma band oscillations in response to sensory stimuli, (McNally et al., 2021), which might limit the effectiveness of these treatments .Indeed, a previous study demonstrated impairments in the ability of hippocampus and visual cortex to entrain to a 40-Hz flickering light stimulus in APP/PS1 and 5XFAD mice (Soula et al., 2023). Based on previous *in vitro* studies in these AD mouse models (Caccavano et al., 2020; Hijazi et al., 2019), abnormal PV neuronal activity could be one of the mechanisms underlying this impairment in evoked gamma oscillations. To investigate this hypothesis, we performed simultaneous recordings of LFP from mPFC and HPC and fiber photometry recordings of calcium activity from hippocampal PV neurons while 40-Hz auditory stimulation (2s-ON, 4s-OFF, 10 kHz at 85 dB, 150 trials) was presented to the mice. We repeated the recording at 3, 4, 5, and 6 months of age in the same mice.

First, we analyzed LFP signals during the 40-Hz auditory stimulation. As expected, 40-Hz auditory stimulation evoked an initial transient broadband cortical response followed by a sustained narrow band response at 40 Hz, known as the auditory steady state response (ASSR) (**Fig. 5A**) (McNally et al., 2021). Although both groups showed response to 40-Hz auditory stimulation (**Fig. 5A**), the steady-state response was weaker in APP/PS1 mice. We focused on the gamma frequency range (35–45 Hz) and computed gamma power based on the LFP data from PFC and HPC during the time of 40-Hz auditory stimulation (0.2 to 4 s from stimulation onset) and compared percentage change from pre-stimulation time (−2 to 0.2 s from stimulation onset) between non-AD and APP/PS1 groups (**Fig. 5B&C**). We found that the ASSR was markedly lower in APP/PS1 mice compared to non-AD mice in the PFC (**Fig. 5B**, Percentage decrease 3mo: 61.15%, 4mo: 62.73%, 5mo: 51.66%, 6mo: 22.71% increase, F1,17= 9.24, p= 0.0074). In the HPC the gamma band response was actually lower than pre-stimulus levels in AD mice (**Fig. 5C**, F1,17= 26.20, p= 0.000086) during the 40-Hz auditory stimulation even at 3 or 4 months of age (post-hoc test 3, 4, 5, 6 months, PFC: p= 0.0082, p= 0.0062, p= 0.086, p= 0.95; HPC: p= 0.13, p= 0.0034, p= 0.011, p= 0.0019).

### Responses of PV neurons in HPC evoked by 40-Hz auditory stimulus were lower in APP/PS1 mice even at 3months old

Next, we looked at fiber photometry data obtained from hippocampal PV neurons. Normalized fluorescent signals aligned at the auditory stimulus onset show sustained PV response in the non-AD group during the auditory stimulation (**Fig. 5D**, solid lines) which are not observed in the APP/PS1 group especially at 3 and 4 months old (**Fig. 5D**, dotted lines). To assess PV neuronal responses to the 40-Hz auditory stimulation at each months, we focused on the time window where sustained PV activity was observed in non-AD animals (0.5–2.5s of auditory stimulus onset, between red dotted lines in **Fig. 5D**) and computed areas under curve and compared them between non-AD and APP/PS1 groups for each month.

Areas under the curve were significantly lower in the APP/PS1 mice (**Fig. 5E**, Percentage decrease 3mo: 86.87%, 4mo: 99.69%, 5mo: 175.45%, 6mo: 75.67%) indicating that responses of PV neurons in hippocampus evoked by 40-Hz auditory stimulus were lower in APP/PS1 mice (F1,17= 8.83, p= 0.0085) as early as at 3 months old (post-hoc test: 3, 4, 5, 6 months, p= 0.028, p= 0.0011, p= 0.030, p= 0.80, respectively) which is about the same time as sleep/wake related brain oscillation changes.

### No impairment in Y-maze spontaneous alternation performance in APP/PS1 mice at least up to 6 months old

To assess the time-course of cognitive decline in APP/PS1 mice, we utilized the Y-maze spontaneous alternation task. Spontaneous Y-maze alternation task measures spatial working-memory and depends on the brain regions including mPFC and HPC (Lalonde, 2002) which are the areas known to begin Aβ accumulation in early AD phase (Palmqvist et al., 2017). Previous studies from other groups demonstrated impairments in this task at around 8 months old in APP/PS1 mice (Da Silva et al., 2016; Hulshof et al., 2022). Consistent with these prior findings, we found no significant difference between non-AD and APP/PS1 mice at any month, suggesting that no apparent cognitive impairment was observed at least up to 6 months old in APP/PS1 mice (**Fig. 6**, F1, 22= 0.015, p= 0.90).

**Fig 6.**
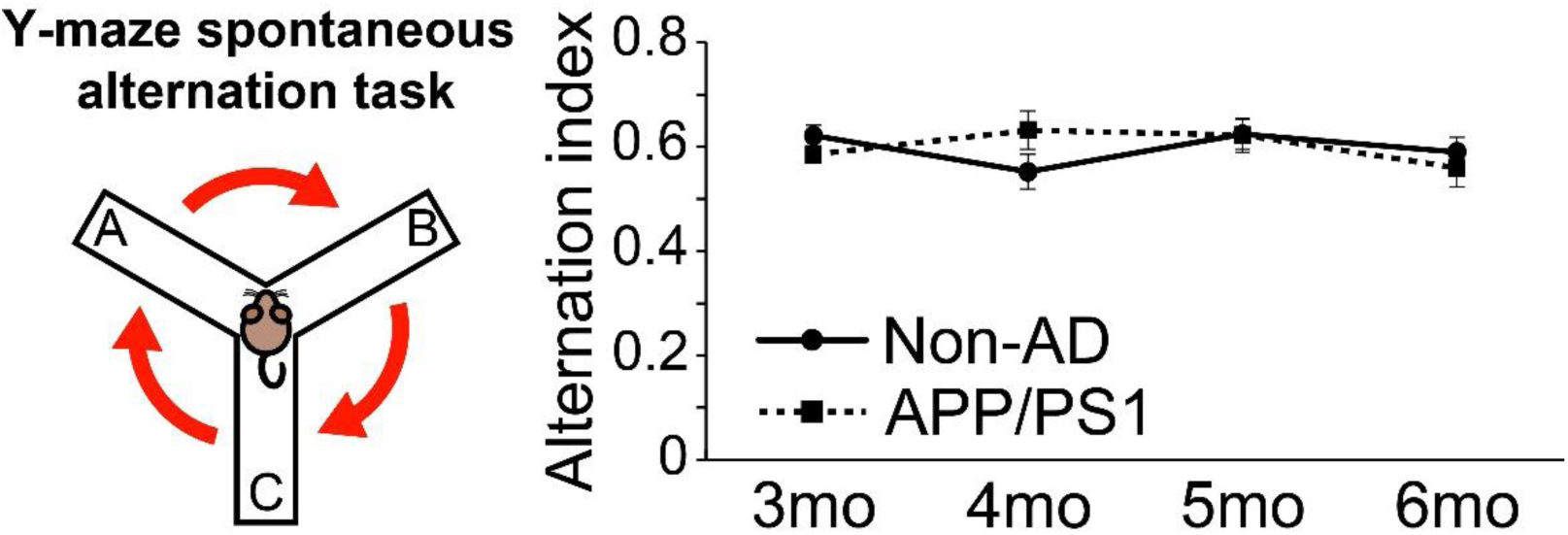
No impairment in Y-maze spontaneous alternation performance in APP/PS1 mice at least up to 6 months old. Cognitive performance was assessed using the Y-maze spontaneous alternation task at each month. No significant difference between Ctr (N=13) and AD (N=11) mice was found at any month, suggesting that no clear cognitive impairment in this task was observed at least up to 6 months old in APP/PS1 mice, in contrast to previous findings in older APP/PS1 mice (Devi & Ohno, 2012; Shukla et al., 2013; Webster et al., 2014).

## DISCUSSION

In the current study, we investigated whether alterations in sleep and hippocampal PV activity occur prior to cognitive deficits in a commonly used amyloid-overexpressing AD mouse model. We performed longitudinal recordings from the same mice *in vivo* at different ages to assess time course of changes in sleep/wake pattern, brain oscillations, activity of hippocampal PV containing neurons, and cognition. This is the first study to report longitudinal *in vivo* recordings of hippocampal PV neurons in AD mouse models along with sleep changes at the early disease phase (3–6 months). Our primary findings are, 1) Band power in slow-wave activity (< 4 Hz) during NREM sleep was markedly lower in the APP/PS1 mice, primarily due to a reduction in the delta (1.5–4 Hz band) linked to the restorative functions of sleep (Huber et al., 2000); 2) the proportion of NREM sleep during the 12h dark (active) phase was significantly lower in APP/PS1 mice compared to non-AD mice, while wakefulness was significantly higher; 3) Band power in the beta frequency range was significantly and consistently higher in the APP/PS1 mice in all sleep/wake states; 4) The 40-Hz auditory-steady state electrical response and the activity of hippocampal PV neurons assessed via fiber photometry recordings during the 40-Hz auditory stimulation was significantly lower in the APP/PS1 mice. These changes were prominent at 3 months and were largely maintained until 6 months of age. Furthermore, reductions in NREM slow-wave activity and enhancements in beta activity in young adults were also observed in another amyloid-overexpressing model, 5XFAD mice. However, no clear cognitive impairment was observed in the Y-maze spontaneous alternation task up until 6 months old in APP/PS1 mice. These results suggest that altered sleep and 40-Hz associated evoked PV neuronal activity are characteristics of earlier stages of AD before apparent cognitive impairment.

### Abnormalities in sleep and brain oscillations in AD

One of the major findings of the current study is very early changes in sleep/wake related brain oscillations in the APP/PS1 mice. Specifically, we observed APP/PS1 mice with lower power in delta range (1.5–4 Hz) and higher power in beta range (15–30 Hz) at early ages before cognitive impairment and brain-wide Aβ accumulation (Garcia-Alloza et al., 2006; Jankowsky et al., 2004) are observed. These findings were accompanied by decreased time spent in NREM sleep and increased time spent in wakefulness in APP/PS1 mice compared to non-AD mice.

In humans, sleep changes become apparent at the preclinical stage of AD and are disrupted further over the course of the disease (Mander et al., 2016). It has been suggested that sleep and AD pathology have a bidirectional relationship, that is, sleep disturbances exacerbate AD pathology including Aβ/tau burden and neuronal dysfunctions, and AD pathology can also worsen sleep disturbances (Y. E. Ju et al., 2014; Mander et al., 2016). Although a causal relationship between sleep disturbances and pathological changes in AD has not yet been proven, finding that sleep abnormalities precede and predict symptoms and pathology adds no weight to the notion that sleep abnormalities are secondary to the disease. Moreover, an increasing number of studies support the link between NREM sleep and the glymphatic system as a key mechanism for clearing metabolic waste products including Aβ and tau in the brain (Harrison et al., 2020; Iliff et al., 2012, 2014; Ishida et al., 2022; Kang et al., 2009b; Xie et al., 2013). The seminal article that launched the glymphatic clearance hypothesis revealed that NREM sleep, particularly slow-wave activity (<4 Hz), facilitates brain clearance in mice (Xie et al., 2013). Slow-wave activity has since been associated with clearance of amyloid precursor protein (Morawska et al., 2016) in mice and Aβ itself in humans (Y. E. S. Ju et al., 2017). However, how slow-wave activity facilitates clearance of toxic waste in the brain is not fully understood. Recent work unveiled part of the mechanism, that is, slow-wave activity recorded by EEG closely precedes large pulses of cerebrospinal fluid (CSF) flowing over the brain (Fultz et al., 2019). Another recent study in mice demonstrated that infraslow norepinephrine oscillations during NREM sleep control tightly linked changes in cerebral blood volume and CSF which promote glymphatic clearance (Hauglund et al., 2025). Our current findings showed that slow-wave activity, especially in the delta range, was already reduced in the APP/PS1 mice at the age when Aβ accumulation was just beginning in the brain (Garcia-Alloza et al., 2006; Jankowsky et al., 2004). This suggests that disrupted slow-wave activity could be one of the key indications of dysfunction of glymphatic system which leads to the progression of AD pathogenesis.

Previous studies have shown altered slow-wave activity in AD mouse models although the timing and directions of alterations vary among the different mouse models and stages of the disease (Katsuki et al., 2022). Consistent with our findings, a previous study reported reduced NREM slow-wave power in frequency band of 0.1–4 Hz was observed in APP/PS1 mice at 8–10 months (Kent et al., 2018), an age-range when more brain-wide Aβ accumulation and cognitive impairments were observed in this model (Garcia-Alloza et al., 2006; Jankowsky et al., 2004). In a study with 5XFAD mice, another Aβ mouse model, the power of the 0.5–4 Hz frequency band was decreased at 6 months compared to 3 months of age or when compared to age-matched non-transgenic mice (Schneider et al., 2014). Our smaller sample of data with 5XFAD mice showed lower slow-wave activity compared to non-AD mice starting around 3 months old which became more significant at 6 months old. The 5XFAD mice are known to exhibit faster progression of AD phonotypes than APP/PS1 mice. In our study, lower power in frequency band of 1.5–4 Hz was consistently found in APP/PS1 mice compared to non-AD mice between 3 and 6 months of age, however, further decline in both slow and delta oscillations could potentially be observed in the later stages of the disease.

Interestingly, we also found that beta power was significantly and consistently higher in APP/PS1 mice. Most reports on sleep oscillations in Alzheimer’s disease so far have primarily focused on lower frequency bands like slow-wave activity that is known to be a hallmark of NREM sleep (Katsuki et al., 2022). However, recent studies in humans and animals reported that enhanced beta activity during sleep is correlated with reduced glymphatic function (Dagum et al., 2025; Hablitz et al., 2019). Hablitz et al. demonstrated that glymphatic function measured as glymphatic tracer influx negatively correlates with EEG beta power (13–20 Hz) while it positively correlates with EEG delta power (1–4 Hz) in the mice under different anesthetic regimens (Hablitz et al., 2019). Consistent with this animal study, more recent human study showed that increased glymphatic function, measured as parenchymal resistance via dynamic impedance spectroscopy, was accompanied by reduced EEG beta power (15–30 Hz) and increased EEG delta power (1–4 Hz) during natural overnight sleep (Dagum et al., 2025).

Beta activity is commonly associated with activity during active wakefulness such as sensory processing and cognitive engagement (Başar-Eroglu et al., 1996; Bouyer et al., 1981; Grønli et al., 2016). Abnormally high beta activity has been linked to central nervous system hyperarousal (Mizrahi-Kliger et al., 2020; Perlis, Merica, et al., 2001; Perlis, Smith, et al., 2001; Riemann et al., 2010; Van Someren, 2021; Verma et al., 2024). Polysomnographic studies in patients with insomnia showed increased beta activity at around sleep onset (Freedman, 1986; Perlis, Merica, et al., 2001; Perlis, Smith, et al., 2001; Strijkstra et al., 2003). These studies indicate that beta activity represents an important functional marker and/or a causal factor in insomnia (Zhao et al., 2021). Our data in APP/PS1 mice show higher beta activity in all states, including in NREM sleep, compared to non-AD mice. Our data from 5XFAD mice also showed higher beta activity compared to non-AD mice. In line with our finding, a previous rodent study in older AD mouse models also showed that beta power (13–20 Hz) was higher in 8–10-month-old APP/PS1 and 12-month-old Tg2576 mice compared to the wildtype mice (Kent et al., 2018). Thus, an increased beta activity could be another unique, early marker of glymphatic dysfunction and sleep disturbances in Alzheimer’s disease.

### Reduced evoked gamma-band activity and altered activities of hippocampal and cortical interneurons in AD

Here we found enhanced spontaneous broad-band gamma and beta power, as well as a reduced evoked auditory 40-Hz steady-state response, a pattern of changes consistent with impaired activity of cortical PV interneurons (McNally et al., 2013, 2021). Alterations in GABA levels and GABAergic interneurons have been reported in both AD patients and animal models of AD (Xu et al., 2020). Reduced PV neurons in cortical regions and hippocampus were reported in AD patients (Brady & Mufson, 1997; Mikkonen et al., 1999; Sanchez-Mejias et al., 2020; Solodkin et al., 1996). In animal models, decreases in the number of cortical and hippocampal PV neurons were reported in 5XFAD mice (Flanigan et al., 2014; Giesers & Wirths, 2020), APP/PS1 mice (Cheng et al., 2020; Popović et al., 2008; Saiz-Sanchez et al., 2012; Takahashi et al., 2010), 3xTg mice (Zallo et al., 2018), and Tg2576 mice (Huh et al., 2016). However, to date longitudinal assessments of the ability of PV neurons to respond to 40-Hz activity *in vivo* were lacking. Previous studies suggested that changes in the activity of cortical and hippocampal PV neurons varies among the models and according to the ages of the animals. *In vitro* work in APP/PS1 mice showed that hippocampal PV neurons are hyperexcitable at an early stage (3–4 months) in disease progression, prior to observable changes in pyramidal neuronal excitability, and that this is linked to abnormal neuronal network activity and memory impairment (Hijazi et al., 2019). Contrary to this finding, another *in vitro* study in 5XFAD mice reported reduced activity of hippocampal PV basket cells especially during sharp-wave ripples in early (∼3 months) amyloid pathology (Caccavano et al., 2020). Hijazi et al. reported that reactivation of PV neurons in 6-month-old APP/PS1 mice had similar benefit on cognition as inhibition of PV neurons in 4-month-old mice. Thus, one possibility is that abnormal activity of hippocampal PV neurons occurs in a biphasic manner, that is, hyperexcitability precedes hypoexcitability (Hijazi et al., 2019). However, in our *in vivo* study in APP/PS1 mice, fiber photometry data recorded during 40-Hz auditory stimulation did not observe a biphasic change in PV activity at least between 3 to 6 months old. The PV neuronal response to the auditory stimulus compared to non-AD mice was consistently lower in the APP/PS1 mice during this experimental period. *In vivo* hypoactivity of PV interneurons is consistent with a previous *in vivo* multiphoton study which recorded spontaneous activity of cortical PV interneurons under isoflurane anesthesia in APP/PS1 mice (Algamal et al., 2022). Reduced spiking and visual tuning of layer 5/6 PV interneurons was also observed in another study before substantial plaque formation and was associated with reduced excitatory synapses and postsynaptic glutamate receptors (Papanikolaou et al., 2025). Thus, even if the intrinsic excitability of PV interneurons is increased *in vitro* (Hijazi et al., 2019; van Adrichem et al., 2025), their activity levels are consistently reduced *in vivo*, likely due to a reduction in excitatory inputs (Papanikolaou et al., 2025).

### Time course of abnormalities in cortical interneurons and brain oscillations in AD

Our study confirmed that disruption in brain oscillations during sleep/wake and abnormal activity of PV neurons both occur early in AD and suggest that they could be potential factors for accelerating AD pathogenesis. However, a question remains: How do all the changes in sleep/wake patterns, relevant brain oscillations, and abnormal neural activity start in AD? A possible explanation is, precisely timed cortical and hippocampal PV activity is critical for the correct generation and nesting of sleep oscillations; Abnormal activity of PV interneurons will lead to disruption of the timing of cortical PV discharge with respect to sleep oscillations and impair calcium dynamics in pyramidal neurons (Averkin et al., 2016; Niethard et al., 2018; Seibt et al., 2017). In fact, chemogenetic excitation of PV interneurons in frontal cortex greatly decreases slow-wave activity during NREM sleep (Funk et al., 2017), indicating that abnormal activity of PV neurons in AD may disrupt cortical excitation/inhibition balance and reduce the slow-wave activity vital for Aβ/tau clearance (Mander et al., 2015; Xie et al., 2013). A study in APP23/PS45 transgenic mice reported disruption in slow-wave activity and the long-range coherence of slow waves in the neocortex, thalamus and hippocampus. They found that the impairments in frequency and long-range coherence of slow waves were rescued by topical applications of GABA_A_ receptor agonist (Busche et al., 2015). Another study reported that cortical GABA levels as well as the expression of GABA_A_ and GABA_B_ receptors were decreased in APP mice at 4 months of age prior to Aβ plaque accumulation along with slow-wave activity disruption (Kastanenka et al., 2017). The same study also showed that the slow-oscillation power was reduced by topical application of GABA_A_ inhibitor onto cortices in healthy control animals. Thus, disrupted cortical excitatory/inhibitory interactions due to alterations of GABA release as well as GABA_A_ and GABA_B_ receptors in the neurons including PV neurons could be implicated in abnormal brain oscillatory activity in AD.

Both genetic and epigenetic factors could be potential causes of abnormalities in PV GABAergic neurons in AD. A recent report in human brain organoids shows apolipoprotein E4 (APOE4), the leading genetic risk factor for Alzheimer’s disease, affects cortical neurodevelopment by reducing cortical neurons and altering GABA-related genes (Meyer-Acosta et al., 2025). Chronic stress could also be a factor for cell loss and degeneration of cortical and hippocampal GABAergic interneurons (Czéh et al., 2015; Serradas et al., 2022; Zaletel et al., 2016). A line of studies show that sleep disruption elevates oxidative stress in PV interneurons in cortex and hippocampus leading to neuronal loss and disrupted neural oscillation (Gao et al., 2025; Harkness et al., 2019). These reports suggest that relationship between altered PV neurons and disrupted sleep could be bidirectional. Thus, interventions to prevent alterations in sleep, related brain oscillations, or PV neuronal activity might all be potentially effective approaches for AD.

### Limitations and future directions

Our data was available only up to 6 months old due to the technical challenges to maintain the quality of electrodes and implants for a prolonged period in the same animals. Thus, further decrease in slow-wave activity or PV neuronal activity within animals could be possible as the disease further progresses beyond 6 months of age. Although the focus here was on PV activity, this same general strategy could be applied to investigate other cellular targets in the future. Funk and colleagues showed that chemogenetic activation of somatostatin (SOM)-positive GABAergic neurons increased slow-wave activity, whereas chemogenetic inhibition of SOM neurons decreased slow-wave activity (Funk et al., 2017). Furthermore, optogenetic excitation of cortical SOM neurons co-expressing neuronal nitric oxide synthase (nNOS) evoked a response that resembles a physiological slow wave (Gerashchenko et al., 2018). Thus, SOM/nNOS interneurons could be another promising therapeutic target to enhance sleep and prevent AD progression. We would also note that although we utilized an Aβ mouse model in this study, it is equally vital to understand how changes in sleep and brain oscillations as well as neuronal activity of different cell types link to tau pathology in the Alzheimer’s disease. Further investigations in these aspects would provide a more complete picture of time course of disease and how we will be able to develop effective interventions.

### Conclusions

Our study found that disruptions in sleep and brain oscillations in APP/PS1 mice occur as early as 3 months old before brain-wide Aβ accumulation and apparent cognitive decline happen. The changes were especially notable in brain oscillations with decreased delta power during NREM sleep and increased beta power during wakefulness and REM sleep. Coinciding with the sleep and oscillation changes, we also found that reduced auditory evoked gamma activity along with lower response of hippocampal PV neurons in APP/PS1 mice as early as 3 months old. This is the first time to confirm altered PV activity *in vivo* longitudinally in APP/PS1 mice along with sleep changes. Results suggest that altered sleep and hippocampal PV neuron activity could be biomarkers of earlier stages of AD before apparent cognitive impairment. Specifically, delta and beta oscillation, as well as PV neurons at early stages of disease could be potential targets for intervention. Furthermore, our results establish APP/PS1 mice as a good model to causally test the relationship between sleep, PV neuronal activity and amyloid-mediated pathology and cognitive deficits and to investigate new experimental strategies for early intervention in AD (Kent et al., 2019; Lee et al., 2020; Ogbeide-Latario et al., 2022).

## ACKNOWLEDGEMENTS

This work was supported by United States Veterans Administration Biomedical Laboratory Research and Development Service Merit Awards I01 BX004673 & I01 RD001372 (REB), I01 BX004500 (JMM) and CDA IK2 BX004905 (DSU), United States National Institute of Health support from NIH K01 AG068366 (FK), RF1 AG061774 (DG), R21 MH125242 (JMM, JTM) and by SURE fellowships from Stonehill College. FK is a Health Science Specialist, JMM and REB are Research Health Scientists at VA Boston Healthcare System, West Roxbury, MA. The contents of this work do not represent the views of the U.S. Department of Veterans Affairs or the United States Government.

## AUTHOR CONTRIBUTIONS

FK: Conceptualization, Methodology, Investigation, Software, Formal analysis, Writing - Original Draft, Visualization, Funding acquisition. JMM: Conceptualization, Methodology, Writing - Review & Editing. DD: Conceptualization, Methodology, Writing - Review & Editing. DSU: Software, Resources, Writing - Review & Editing. AT: Investigation, Writing - Review & Editing. JGM: Investigation, Writing - Review & Editing. JTM: Methodology, Resources, Writing - Review & Editing. REB: Supervision, Conceptualization, Methodology, Writing - Review & Editing.

---

**Supplemental Fig 1.**
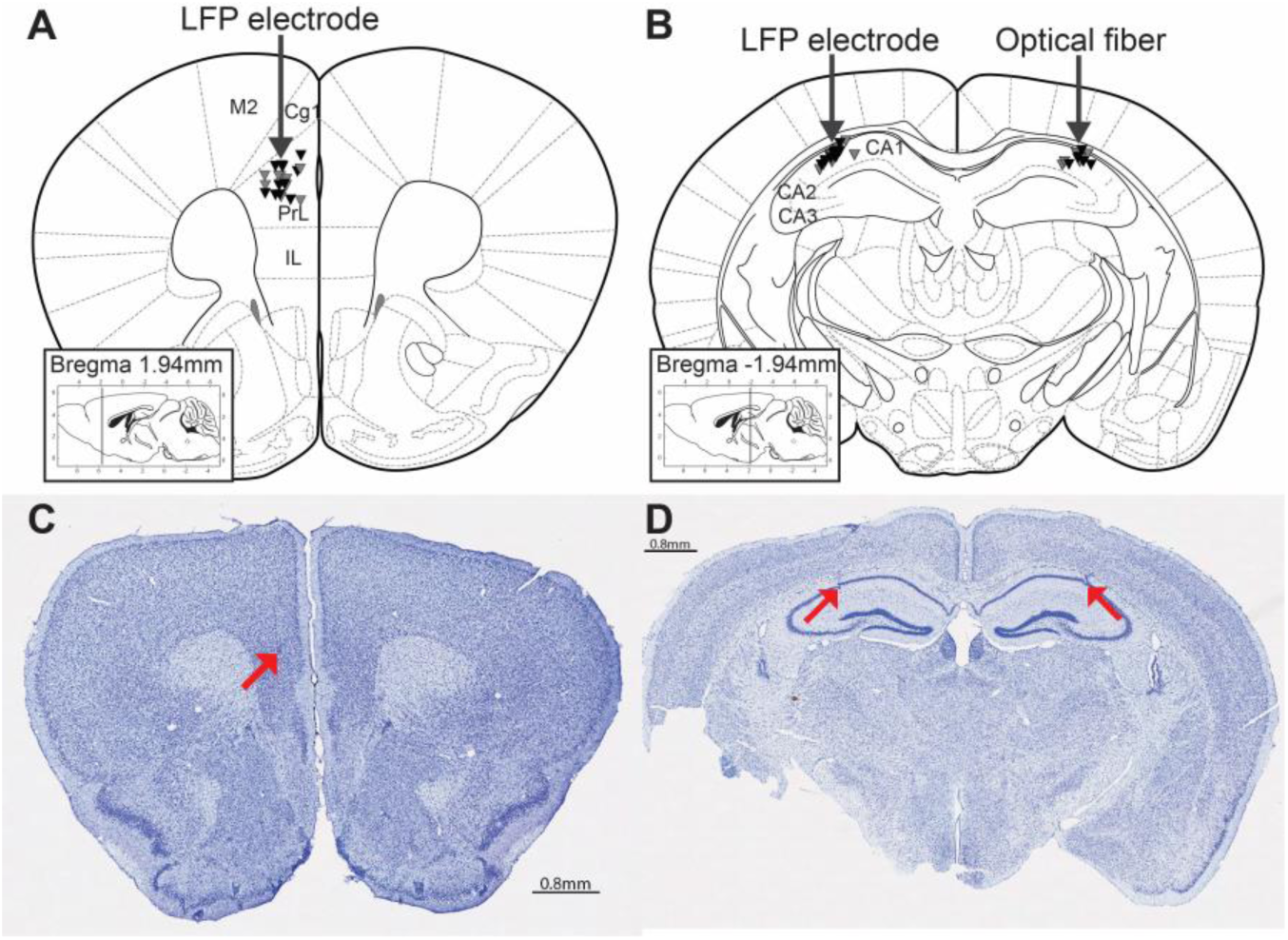
Histological verification of electrode/optical fiber targeting and GCaMP transduction. **(A&B)** Locations of LFP electrodes and optical fibers were confirmed in PFC (**A**) and hippocampal CA1 (**B**) with cresyl violet staining. **(C&D)** Representative images of PFC (**C**) and hippocampal CA1 (**D**) with cresyl violet staining showing the examples of LFP electrodes and optical fiber traces (red arrows).

**Supplemental Fig 2.**
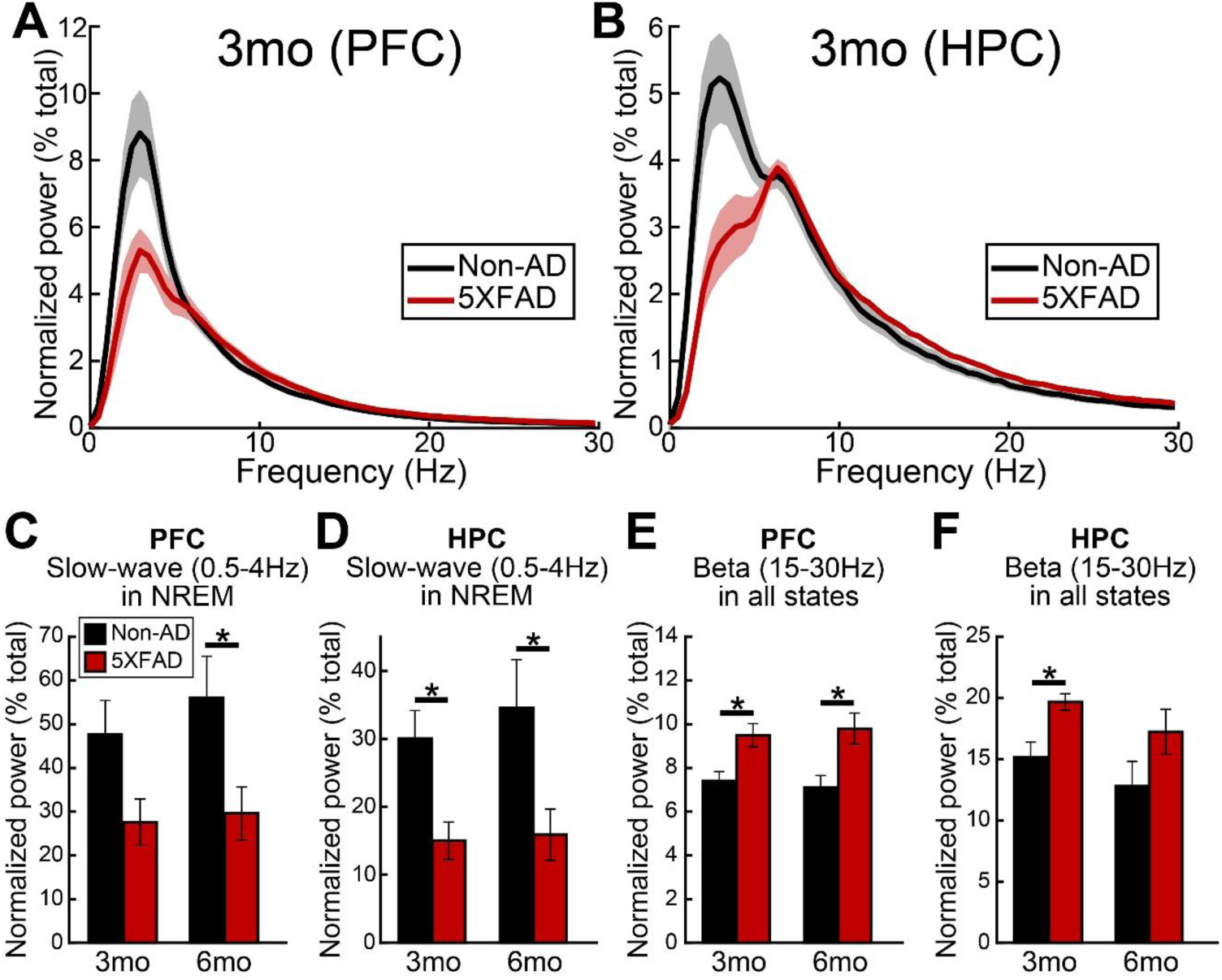
Lower slow-wave activity and higher beta activity were observed in 5XFAD mice compared to the wild-type mice as early as 3 months old. **(A&B)** Averaged power spectral density (PSD) plots from PFC (A) and HPC (B) LFP recordings showed lower power around the slow-wave activity range (0.5–4Hz) in 5XFAD mice (N=6) compared to non-AD mice (N=7) at 3 months of age. **(C&D)** Slow-wave activity (0.5–4Hz) was reduced during NREM sleep in 5XFAD mice (N=6) as early as 3 months old compared to non-AD mice (N=7) in (**C**) PFC and (**D**) HPC. **(E&F)** Beta power (15–30Hz) was higher in 5XFAD mice in (**E**) PFC and (**F**) HPC when data from all states are analyzed together. Data during NREM sleep of 24-h recordings were used for the analysis. Shaded areas and error bars represent SEM. Repeated measures ANOVAs with post-hoc Tukey-Kramer test was performed (*p<0.05).

**Supplemental Fig 3.**
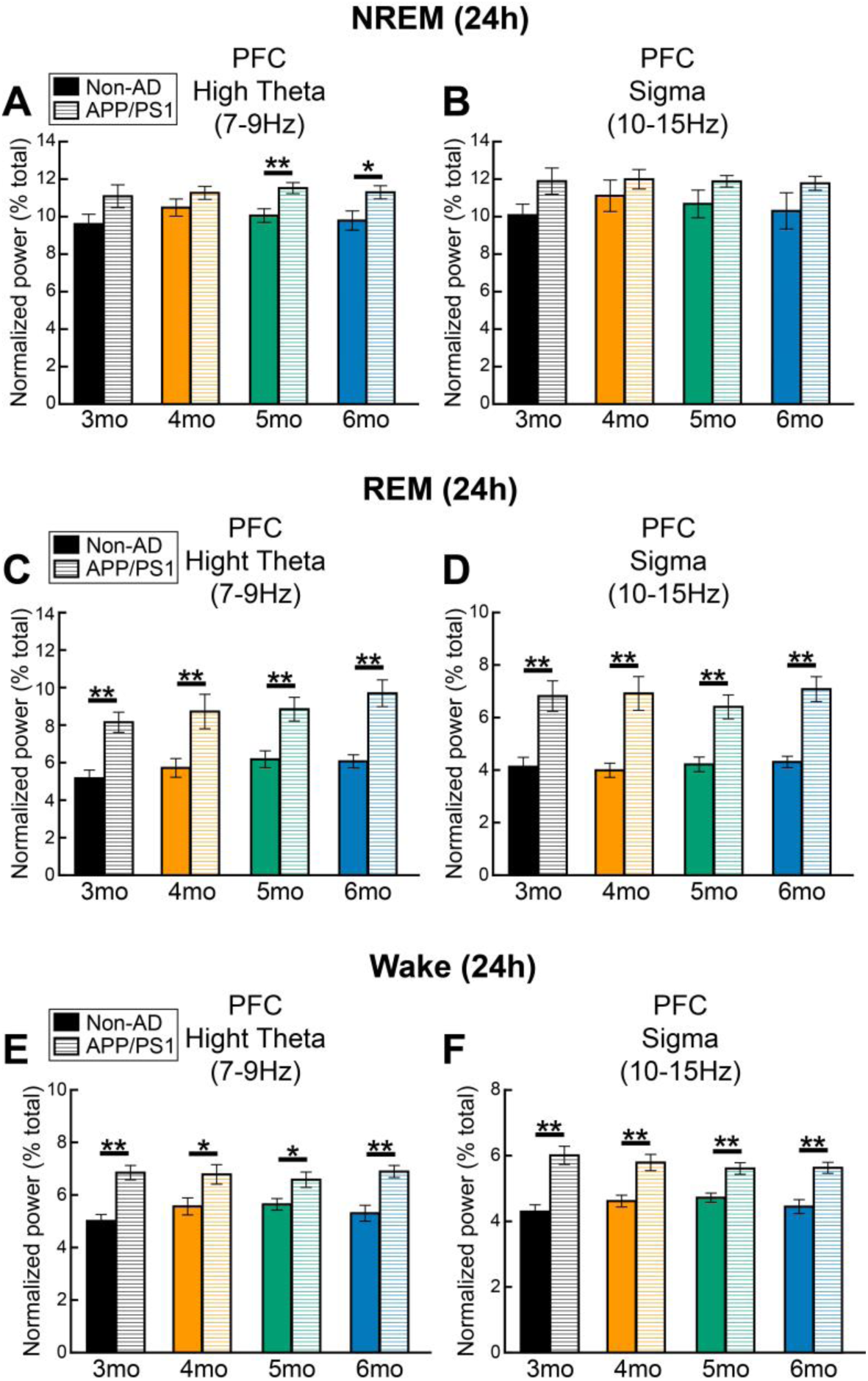
Theta and sigma power in PFC was higher in APP/PS1 mice as early as 3 months old but differences were state dependent. **(A, C, E)** Spectral power around high theta range (7–9 Hz) was significantly higher in APP/PS1 mice (N=11) in PFC especially during REM sleep (**C**) and wakefulness (**E**) compared to non-AD mice (N=13). This difference was observed from 3 to 6 months old. **(B, D, F)** Sigma power (10–15 Hz) was also higher in APP/PS1 mice compared to the wild type mice during REM sleep (**D**) and wakefulness (**F**), but not during NREM sleep (**B**). Data from both light and dark phase were included. Error bars represent SEM. Repeated measures ANOVAs with post-hoc tukey-kramer test was performed (*p<0.05, **p<0.01).

